# Meta3C analysis of a mouse gut microbiome

**DOI:** 10.1101/034793

**Authors:** Martial Marbouty, Lyam Baudry, Axel Cournac, Romain Koszul

## Abstract

Microbial populations as well as they biochemical activities are important components of environmental ecosystems, including the human microbiome. Deciphering the genomic content of these complex mixes of species is an important challenge but is essential to fully understand the regulation of their ecological balance. Here we apply meta3C, an experimental and computational approach that exploits the physical contacts between chromosomes to characterize large genomic regions of bacterial species mixed together, on a truly complex ecosystem: the mouse gut microbiota. Meta3C, which was initially described and applied onto controlled mixes of microorganisms, allowed the *de novo* assembly and scaffolding of numerous bacteria present into this natural mix. Importantly, the scaffolds analyzed exhibit the structural properties expected from typical bacterial chromosomes. Meta3C therefore paves the way to the in-depth analysis of genomic structuration of complex populations.

## Introduction

The collisions experienced by DNA segments belonging to one or several chromosomes and constrained within a cellular compartment generate unique physical 3D signatures that can be exploited to improve genomic analyses (Flot et al., 2014; Marbouty and Koszul, 2015). Such “contact genomics” approaches have recently been used to improve haplotype phasing (Selvaraj et al., 2013) and chromosome scaffolding (Burton et al., 2013; Kaplan and Dekker, 2013; Marie-Nelly et al., 2014a) as well as to annotate genomes (Marie-Nelly et al., 2014b), and have shown promising perspectives in the field of metagenomics. Using controlled mixes of microorganisms, we (Marbouty et al., 2014) and others (Beitel et al., 2014; Burton et al., 2014) showed that, indeed, the genomes of species present into a mix could be partially characterized through a clustering analysis of DNA contacts quantified using chromosome conformation capture (3C) (Dekker et al., 2002). The original proof-of-concept meta3C analysis was performed on both controlled mixes and onto a semi-natural unknown mix of microorganisms (Marbouty et al., 2014). The blind analysis of the meta3C paired-end reads gave promising results: briefly, the reads were used to generate a *de novo* assembly, and the paired-end information was exploited to pool the resulting contigs into communities through a partition detection algorithm (Marbouty and Koszul, 2015; Marbouty et al., 2014). Contigs within each community were then scaffolded into large chromosomal regions using our homemade program GRAAL (Marie-Nelly et al., 2014a). The meta3C on the unknown semi-natural environmental sample showed that large amounts of DNA segments could be associated as part of an intricate network of 3D interactions and called for the analysis of truly natural, complex ecosystems. Here, we show that meta3C indeed fulfills its promises when it comes to a truly natural, complex microbial community: the mouse gut microbiome. From a set of two meta3C libraries generated with two frequent cutter enzymes recognizing either GC or AT-rich sequences, we were able to characterize *de novo* near-complete or important portions of the genome sequences from species present within the mix. Meta3C therefore paves the way to the full characterization of gene content of bacteria present in complex communities, including its dynamic changes resulting from transfers of genes, plasmids, or phages.

## Results and discussion

We collected feces of healthy control mouse (C57BL/6) from the Institut Pasteur animal facility and processed them using two meta3C protocols that solely differed by the restriction enzyme, HpaII (CCGG) or MluCI (AATT) (Figure 1A; Marbouty et al., 2014). As discussed in the perspectives of our original publication, we expected that using two enzymes differing in the GC content of the corresponding restriction sites (RS) would improve the equal coverage of contacts for GC-rich and AT-rich genomes. The two libraries were paired-end (PE) sequenced on an Illumina NextSeq machine (2x65bp), with 114 and 71 million PE reads for libraries HpaII and MluCI, respectively. Reads from both libraries were then pooled and assembled into contigs using the IDBA-UD program (Peng et al., 2012), resulting in 373,363 contigs (cumulated size: 580 Mb, N50: 3,783 pb, max size: 490Kb, mean size: 1,402bp). This assembly was analyzed at the taxonomic level using the MG-RAST pipeline (Meyer et al., 2008). As expected from a gut metagenome, the major clades in the sample were Clostridia (65%) and Bacteroidetes (15%) (Figure 1B; Langille et al., 2014). Interestingly, a similar analysis performed on the DNA sequences using the Kraken program (Wood and Salzberg, 2014) confirmed these results but also showed that the major part (ca. 80%) of the DNA sequences could not be attributed to a sequenced organism (data not shown).

**Figure 1.**
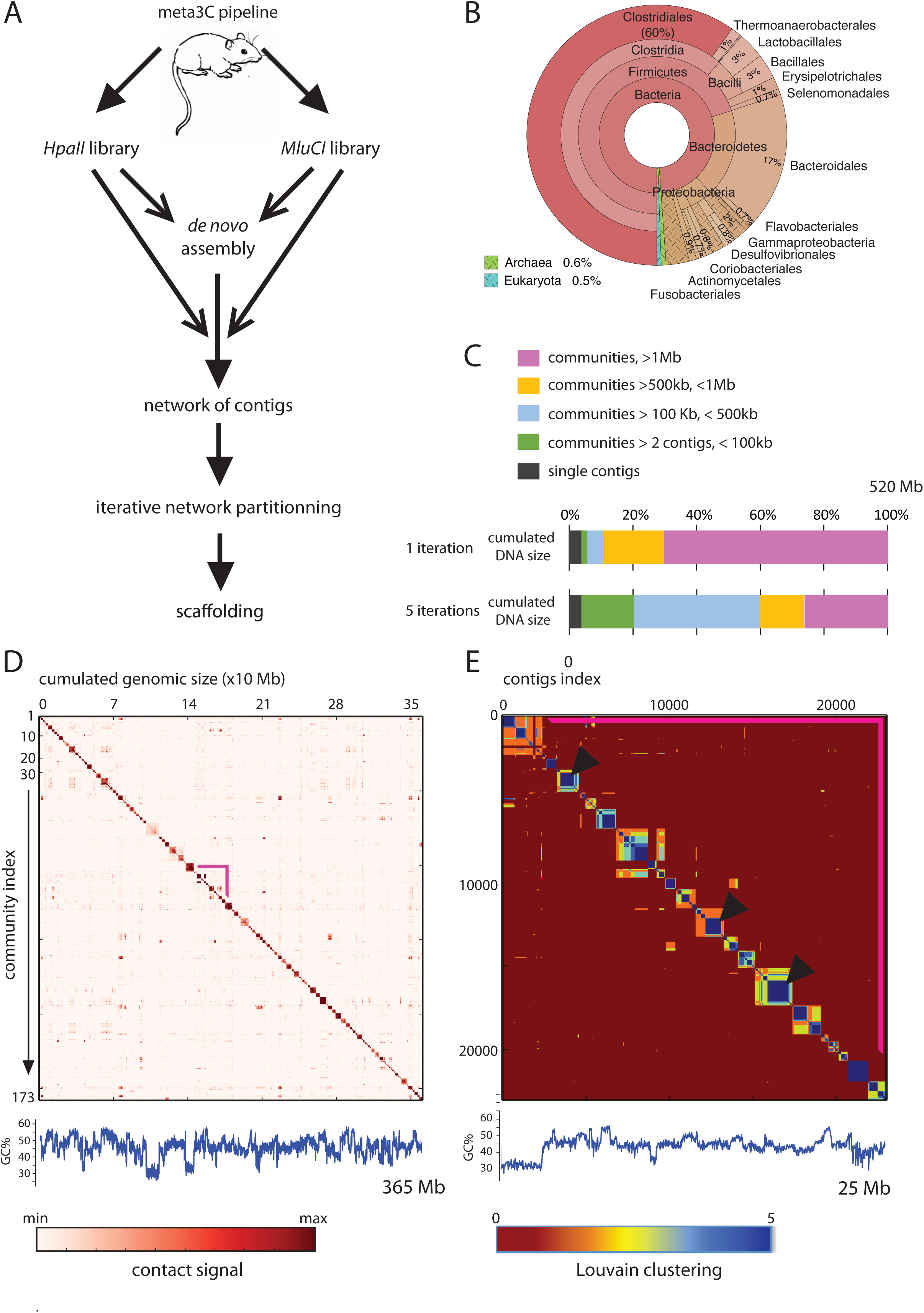
meta3C analysis of the mice gut microbiome. **A**, Flowchart representing the computational analysis steps of a meta3C experiment. First, the reads from two sequenced meta3C libraries are assembled into contigs. The meta3C contact information contained in the 2 datasets is then used to generate a contact network between all the contigs. The Louvain algorithm is then applied iteratively to segment the global network into smaller overlapping communities. Ultimately, the contigs present in each overlapping community are scaffolded using GRAAL (Marie-Nelly et al., 2014a). **B**, MG-RAST taxonomy analysis of the contigs generated from the *de novo* assembly step. **C**, Characteristics of the communities obtained after one and five iterations of the Louvain algorithm. **D**, Contact map of the 173 communities larger than 1Mb obtained after one segmentation using the Louvain algorithm (1 vector = 200kb). The x and y axis are labelled with the cumulated DNA size and the index of the community, respectively. The color scale is indicated below the map. Pink bars: region shown in panel **E**. Bottom: GC% of the contigs. **E**, Overlapping communities. Here the color code reflects how many times two contigs cluster together after five iteration of the Louvain algorithm (from one, orange, to five, dark blue). The region represented is a magnification of the region indicated with pink bars in panel **D**. Bottom: GC% of the contigs. Dark triangles points to core communities containing more than 1 Mb of DNA sequences (see Supplementary Figure 1.1.A)

The contigs were then split into 1-kb fragments. This step had two objectives: first, to alleviate the impact of potential misassemblies arising during the assembly step; second, to normalize the contact signal with respect to the influence of contig size on their representation. Contigs under 500bp were discarded, leading to a global set of 553,000 contigs corresponding to an assembly of 520Mb (Figure 1C). Importantly, small contigs discarded at this step can still be exploited later during the analysis (see below). Reads from both meta3C libraries were fused and aligned against these contigs, generating a large network (553,000 nodes, 46 million edges). In order to identify subnetworks likely to correspond to DNA molecules sharing the same cellular compartment (Marbouty et al., 2014), this network was segmented into 47,140 communities (in a network analysis sense) using the Louvain clustering algorithm (Blondel et al., 2008; Material and Methods). 42,613 communities contained only one contig (~4% of the total DNA), while 4,527 contained two contigs or more with a cumulated DNA content ranging from 2kb to 6Mb (Figure 1C). Overall, 465 Mb (~90%) were contained within 317 communities ranging in size from 0.5Mb to 6Mb. The size of the DNA content of the communities identified in the meta3C data are consistent with the range of genome sizes commonly found among bacteria (from 500 Kb to 10 Mb; Islas et al., 2004). The 173 communities containing 1Mb or more were plotted as a contact map with fixed-size bins (Figure 1D). Interestingly, the strong influence of the choice of the restriction enzyme on the representation of the contigs appeared clearly when the same contact map was binned as a function of a fixed number of restriction fragments for each enzyme (Supplementary Figure 1.1). Under this representation the size occupied by a community is directly correlated with its average GC content: contigs presenting a high GC content are frequently split by an enzyme recognizing a GC-rich RS, while an enzyme recognizing an AT-rich RS will restrain the visibility of these contigs during the 3C experiment (Cournac et al., 2015; Imakaev et al., 2012). For each enzyme, the normalized percentage of contacts exploited to generate one community was computed (Supplementary Figure 1.1C). This experiment illustrates the interest of combining two different restriction enzymes to perform a meta3C experiment.

By design, the Louvain algorithm does not attribute a node to several communities. Interestingly, when the segmentation was performed twice on the same network some nodes were attributed to distinct partitions, suggesting these communities shared the sequences represented by these nodes. In order to estimate the likelihood that a contig equally connected to numerous subnetwork will be attributed to its appropriate community/ies, Louvain clustering was run five times independently on the same dataset, exploiting the inherent non-deterministic nature of this algorithm (Figure 1E). This process allowed us to characterize core communities, i.e. sets of contigs that systematically clustered together during the five iterations (Figure 1C and Figure 1E – blue squares). 76 core communities encompassing more than 1Mb were detected (Figure 1E – blue squares pointed by black arrows and Supplementary Figure 2.1.A) and 126 encompassing more than 500 Kb. Each core community above 500 Kb was then extended to all contigs clustering at least once during the 5 iterations with that core, to generate “overlapping communities” (Figure 2 and Supplementary Figure 2.1.B). Among those overlapping communities, some contained several cores encompassing more than 500 Kb and could possibly reflect multipartite genomes (Supplementary Figure 2.1.B). Importantly, this approach allows taking into consideration conserved/repeated sequences or episomes shared by several species (i.e. different core communities above 500 Kb) and, consequently, significantly increases the size of the recovered overlapping communities.

**Figure 2.**
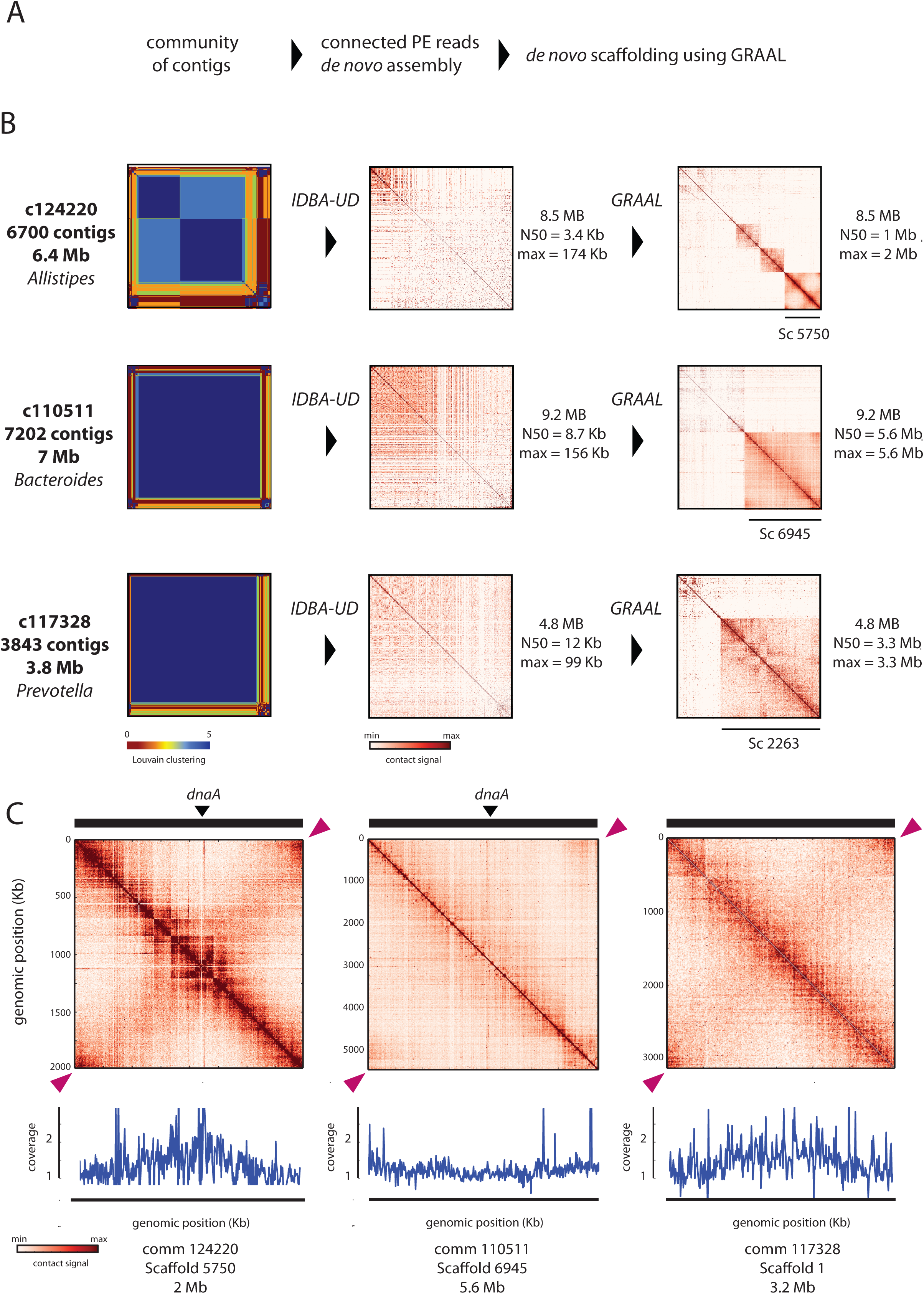
Scaffolding of bacterial genomes in the mice gut microbiome. **A**, Flowchart representing the steps performed to assemble overlapping communities. Contigs from a core community (left) were retrieved and used to build a genome index. The PE reads from the two meta3C libraries were mapped against this index and all the PE reads exhibiting at least one hit were retained for an assembly using IDBA-UD (middle). The scaffolding of the resulting contigs was performed using GRAAL (right). **B**, Examples of the contact map generated at each step of the process for 3 communities. From left to right: overlapping communities; contact map of the contigs obtained from the *de novo* assembly step; and contact map obtained after scaffolding with GRAAL. Quantifications of the assemblies are indicated. **C**, Contact map of the three main scaffolds obtain for the communities shown in panel **B**, right, after GRAAL processing (10 kb bins). Genomic positions are indicated on each map. The variation in coverage is indicated under each map, supposedly reflecting the growth rate of corresponding bacteria. Pink triangles point at the signal indicative of the circular nature of the scaffold, consistent with bacterial circular chromosome.

In order to further investigate the significance of these overlapping communities and assess their genomic content, a *de novo* assembly step was performed for six of them as follow (Figure 2A and Supplementary Figure 2.1). First, all the meta3C PE reads from the original libraries were aligned against the set of contigs of each community (parameters: --local --fast). All read pairs for which at least one read aligned against one of the contigs were retained for a *de novo* assembly using IDBA-UD (N50 from 1.3 Kb to 24 Kb; Table 1). For each community the GRAAL scaffolding program was then run for 10 iterations (Marie-Nelly et al., 2014a) (Table 1). Remarkably, GRAAL assembly resulted in an impressive increase in the N50 which jumped to the megabase scale for 5 of the 6 communities (Figure 2B) (for the sixth community, one large scaffold was retrieved (6.7 Mb) that did not cover half of the assembly).

Each large scaffold obtained at this step was then individually reanalyzed in order to check for DNA fiber continuity. Remarkably, the pattern displayed by the contact maps of these scaffolds appeared highly consistent with published bacterial genomic contact maps (Le et al., 2013; Marbouty et al., 2014, 2015). First, the main diagonal displayed an enrichment in local contacts, resulting from the fact that neighboring DNA regions interact more often together than distant ones. Moreover, a circular signal for 3 of these large scaffolds was clearly apparent on the contact maps, indicating that most of the circular chromosome was retrieved (pink arrowheads, Figure 2C). Interestingly, secondary features characteristic of chromosome metabolism were also visible in some of the maps, such as cohesion of replichores initiated at the origin of replication (Figure 2C; Marbouty et al., 2015). To support this observation, a *dnaA* gene homolog was identified at the crossing between this secondary and the main diagonal (no homologous could be find for the scaffold retrieved from the community 117328). *dnaA* is found at the origin of replication in most bacteria, and its presence at the edge of the secondary diagonal is highly consistent with recent analysis describing the role of the origin of replication during the cell cycle of *Bacillus subtilis* in chromosome folding (Marbouty et al., 2015). Other species such as *Escherichia coli* do not display such contacts, and their genomic contact maps are more similar to the one observed for community 110511 (Marbouty et al., 2014; MM and RK unpublished). Alternatively, the read coverage suggest this species is not dividing, which may also account for the loss of cohesion between replichores (Marbouty et al., 2015; unpublished data). Among the six overlapping communities studied, two were made of several large core communities. However, the reconstructed matrix did not display clear signal indicating a possible multipartite genome. Deeper deciphering of the structure of these particular overlapping communities will be needed to understand their full composition.

It is worth noting that this approach does not require multiple experiments: a single meta3C library generated with a single restriction enzyme will still bring an important amount of genomic information. Also, we noticed that overlapping communities sometimes contain well-individualized scaffolds that are connected through tenuous contacts. One interpretation is that some species or strains contain similar DNA sequences, including plasmids and phages, whose contact signal artificially bridges the genomes of different species. Although we searched for confirmation of this hypothesis within the reads, we could not confirm nor infirm this hypothesis, most likely because of lack of sequencing depth. More analyses and studies should clarify these links in the future.

Overall, the first meta3C experiment performed on a truly complex natural microbiome, brings promising perspectives to this field by highlighting the power of contact genomics approaches to tackle ecological microbial complexity.

## Material and Methods

### Construction of a the meta3C library of a mouse gut microbiota

Feces from C57BL6 male moss were recovered and immediately suspended in 30 mL of TE buffer 1X supplemented with 3% of fresh formaldehyde. Fixation proceeded for 1h under gentle agitation. 10 mL of glycine 2.5M was added to the tube and the quenching was performed for 20 min. The resulting material was washed and recovered by centrifugation and store at −80°c until use. Meta3C libraries were then prepared as described in Marbouty et al. (2014).

### Illumina sequencing

Illumina sequencing was performed as described in Marbouty et al. (2014).

### Genome assembly

Reads containing undetermined bases were removed before the assembly step to retain only good quality reads. *De novo* assemblies were then performed using the program IDBA-UD (Peng et al., 2012) without pre-correction option and default parameters.

### Binning of contigs using 3C contact data

An approach similar to the one described in Marbouty et al. (2014), based on the Louvain method (Blondel et al., 2008), was used to group the different contigs into communities reflecting the different genomes present in the sequenced mixtures. Before applying the algorithm, contigs were divided into 1-kb chunks corresponding to several nodes in the graph. In order to improve the reliability and stability of the clustering, five iterations of the Louvain algorithm were independently run on the dataset, using its non-deterministic heuristics to determine how often contigs clustered with each other. A set of six such clusters comprising large overlapping communities of contigs was selected for further reassembly by GRAAL.

### GRAAL scaffolding of overlapping communities

The GRAAL algorithm was initialized with each set of contigs and an iterative assembly was performed for 10 iterations as described in Marie-Nelly et al. (2014a). The principles of GRAAL scaffolding are described in Marbouty et al. (2014) and Marie-Nelly et al. (2014a). Briefly, after initialization with a set of contigs and the corresponding 3C contact data, GRAAL splits these contigs into smaller fragments that are tested several times to refine their position relative to each other. We used GRAAL to reassemble the contigs contained in each overlapping community previously selected from the Louvain clustering. Table 1 recapitulates the outcome of this scaffolding step, reflected by a strong increase in the N50 parameter and the generation of large (> 1 Mb) scaffolds exhibiting the properties of bacterial genomes.

## Acknowledgements

We thank Julien Mozziconacci for helpful suggestions, Jean-Francois Flot for performing the original IDBA-UD assembly and Kraken analysis as well as for fruitful discussions, and Thierry Pedron for kindly providing us with the mouse feces. This research was supported by funding to R.K. from the European Research Council under the 7th Framework Program (FP7/2007-2013)/ERC grant agreement 260822.

All authors have seen and approved the manuscript.

## Figure supplement legends

**Figure 1 – figure supplement 1.**
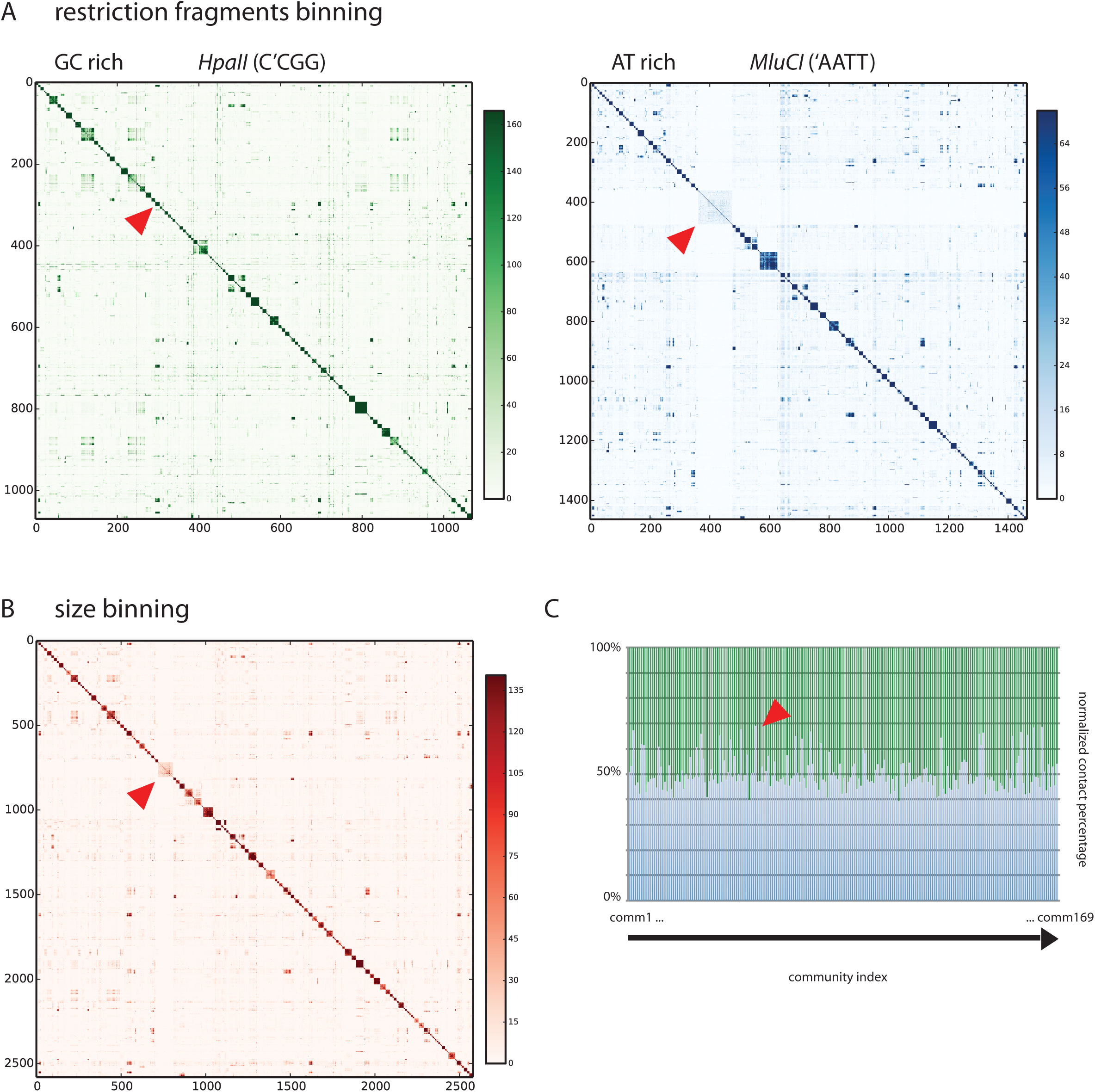
**A**, Contact map of the 173 communities above 1 Mb obtained after 1 iteration of the Louvain algorithm using either *HpaII* or *MluCI* as restriction enzyme. Bins of the two maps correspond to a fixed number of restriction fragments. This representation clearly shows the different response of communities to the two enzymes as indicated by the red triangles. **B**, Combined contact map binned at the Kb level. **C**, Cumulative histogram of normalized contact due to each enzyme (green – *MluCI;* blue – *HpaII)* for the 173 communities.

**Figure 2 – figure supplement 1.**
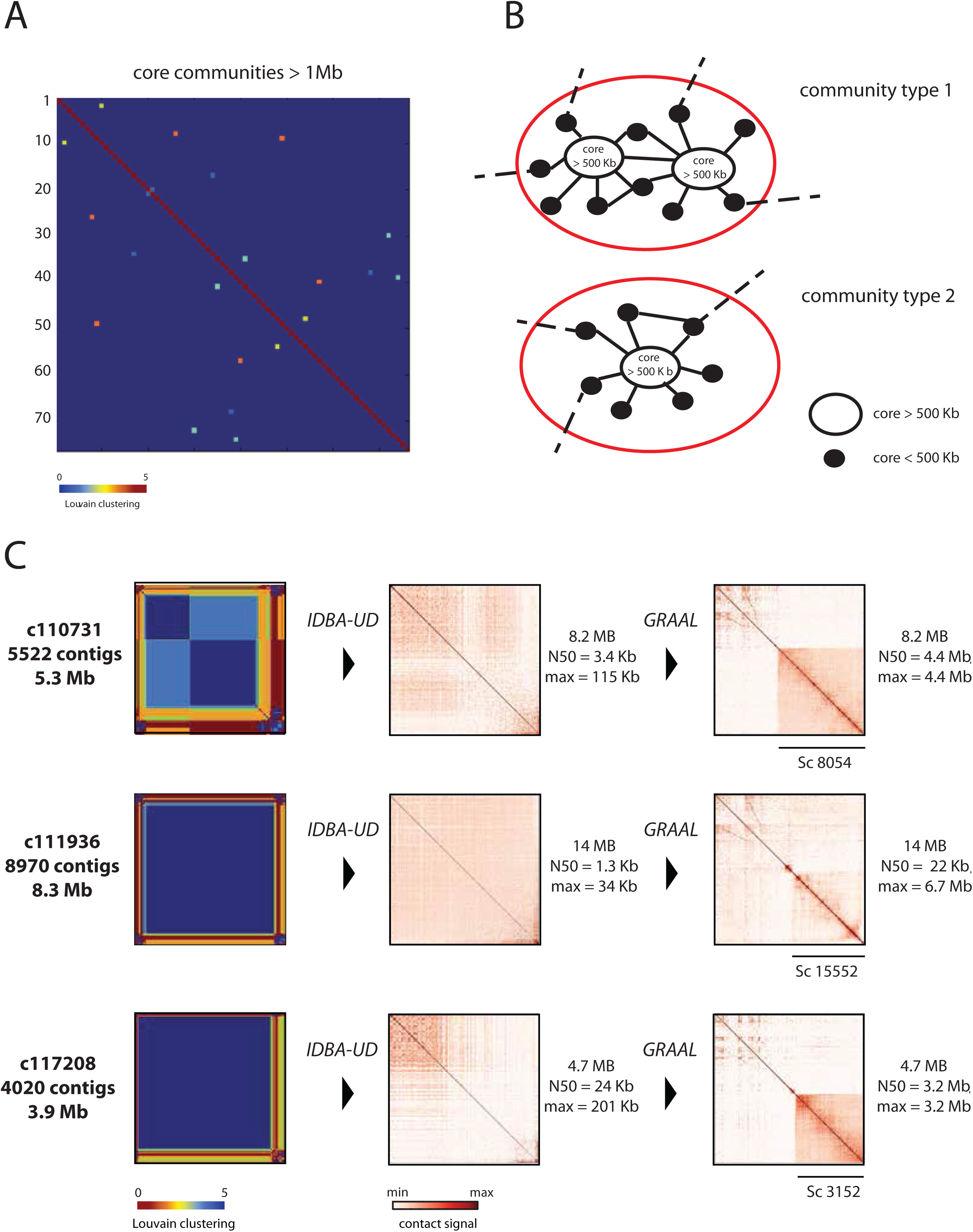
**A**, Map of the 76 core communities containing each more than 1 Mb obtained after the five Louvain iterations. Color code range from 0 (blue) to 5 (red). **B**, Illustration of the process applied to recover overlapping communities. Different type of overlapping communities are illustrated: one that contain only one core above 500 Kb and one that contain two cores above 500 Kb. **C**, Analysis of three additional overlapping communities (see Figure 2B).

**Table 1.**

| Community index | Clustering data |  | IDBA-UD Assembly |  | GRAAL scaffolding |  |
| --- | --- | --- | --- | --- | --- | --- |
| 110511 | # contigs | 7202 | size | 9.2 Mb | size | 9.2 Mb |
|  | cumulated size | 7 Mb | contigs | 7723 | scaffolds | 6945 |
|  | *HapII* | 13 millions | **N50** | **8.7 Kb** | **N50** | **5.6 Mb** |
|  | *MluCI* | 5.5 millions | **max** | **156 Kb** | **max** | **5.6 Mb** |
| 111936 | # contigs | 8970 | size | 14 Mb | size | 14 Mb |
|  | cumulated size | 8.3 Mb | contigs | 21579 | scaffolds | 1552 |
|  | *HapII* | 8 millions | **N50** | **1.3 Kb** | **N50** | **22 Kb** |
|  | *MluCI* | 2.8 millions | **max** | **34 Kb** | **max** | **6.7 Mb** |
| 124220 | # contigs | 6700 | size | 8.5 Mb | size | 9.2 Mb |
|  | cumulated size | 6.4 Mb | contigs | 7968 | scaffolds | 5750 |
|  | *HapII* | 3.8 millions | **N50** | **3.4** | **N50** | **1.1 Mb** |
|  | *MluCI* | 1.7 millions | **max** | **174 Kb** | **max** | **2.06 Mb** |
| 110731 | # contigs | 5522 | size | 8.2 Mb | size | 8.2 Mb |
|  | cumulated size | 5.3 MB | contigs | 9538 | scaffolds | 8054 |
|  | *HapII* | 2.7 millions | **N50** | **3.4 Kb** | **N50** | **4.4 Mb** |
|  | *MluCI* | 1 millions | **max** | **115 Kb** | **max** | **4.4 Mb** |
| 117328 | # contigs | 3843 | size | 4.8 Mb | size | 4.8 Mb |
|  | cumulated size | 3.8 Mb | contigs | 3680 | scaffolds | 2263 |
|  | *HapII* | 1 millions | **N50** | **12 Kb** | **N50** | **3.3 Mb** |
|  | *MluCI* | 0.6 millions | **max** | **99 Kb** | **max** | **3.3 Mb** |
| 117208 | # contigs | 4020 | size | 4.7 MB | size | 4.7 MB |
|  | cumulated size | 3.9 Mb | contigs | 3422 | scaffolds | 3152 |
|  | *HapII* | 2.1 millions | **N50** | **24 Kb** | **N50** | **3.2 Mb** |
|  | *MluCI* | 0.9 millions | **max** | **201 Kb** | **max** | **3.2 Mb** |

## References

Beitel, C.W., Froenicke, L., Lang, J.M., Korf, I.F., Michelmore, R.W., Eisen, J.A., and Darling, A.E. (2014). Strain- and plasmid-level deconvolution of a synthetic metagenome by sequencing proximity ligation products. PeerJ 2, e415.

Blondel, V.D., Guillaume, J.-L., Lambiotte, R., and Lefebvre, E. (2008). Fast unfolding of communities in large networks. J. Stat. Mech. Theory Exp. 2008, P10008.

Burton, J.N., Adey, A., Patwardhan, R.P., Qiu, R., Kitzman, J.O., and Shendure, J. (2013). Chromosome-scale scaffolding of de novo genome assemblies based on chromatin interactions. Nat. Biotechnol. 31, 1119–1125.

Burton, J.N., Liachko, I., Dunham, M.J., and Shendure, J. (2014). Species-level deconvolution of metagenome assemblies with Hi-C-based contact probability maps. G3: Genes|Genomes|Genetics 4, 1339–1346.

Cournac, A., Marbouty, M., Mozziconacci, J., and Koszul, R. (2015). Generation and analysis of chromosomal contact maps of yeast species. Methods Mol. Biol.

Dekker, J., Rippe, K., Dekker, M., and Kleckner, N. (2002). Capturing chromosome conformation. Science 295, 1306–1311.

Flot, J.-F., Marie-Nelly, H., and Koszul, R. (2015). Contact genomics: scaffolding and phasing (meta)genomes using chromosome 3D physical signatures. FEBS Lett. 589, 2966–2974

Imakaev, M., Fudenberg, G., McCord, R.P., Naumova, N., Goloborodko, A., Lajoie, B.R., Dekker, J., and Mirny, L.A. (2012). Iterative correction of Hi-C data reveals hallmarks of chromosome organization. Nat. Methods 9, 999–1003.

Islas, S., Becerra, A., Luisi, P.L., and Lazcano, A. (2004). Comparative genomics and the gene complement of a minimal cell. Orig. Life Evol. Biosphere J. Int. Soc. Study Orig. Life 34, 243–256.

Kaplan, N., and Dekker, J. (2013). High-throughput genome scaffolding from *in vivo* DNA interaction frequency. Nat. Biotechnol. 31, 1143–1147.

Langille, M.G., Meehan, C.J., Koenig, J.E., Dhanani, A.S., Rose, R.A., Howlett, S.E., and Beiko, R.G. (2014). Microbial shifts in the aging mouse gut. Microbiome 2, 1.

Le, T.B.K., Imakaev, M.V., Mirny, L.A., and Laub, M.T. (2013). High-resolution mapping of the spatial organization of a bacterial chromosome. Science 342, 731–734.

Marbouty, M., and Koszul, R. (2015). Metagenome analysis exploiting high-throughput Chromosome Conformation Capture (3C) data. Trends Genet., in press

Marbouty, M., Cournac, A., Flot, J.-F., Marie-Nelly, H., Mozziconacci, J., and Koszul, R. (2014). Metagenomic chromosome conformation capture (meta3C) unveils the diversity of chromosome organization in microorganisms. eLife 3, e03318.

Marbouty, M., Le Gall, A., Cattoni, D.I., Cournac, A., Koh, A., Fiche, J.-B., Mozziconacci, J., Murray, H., Koszul, R., and Nollmann, M. (2015). Condensin- and replication-mediated bacterial chromosome folding and origin condensation revealed by Hi-C and super-resolution imaging. Mol. Cell 59, 588–602.

Marie-Nelly, H., Marbouty, M., Cournac, A., Flot, J.-F., Liti, G., Parodi, D.P., Syan, S., Guillén, N., Margeot, A., Zimmer, C., et al. (2014a). High-quality genome (re)assembly using chromosomal contact data. Nat. Commun. 5, 5695.

Marie-Nelly, H., Marbouty, M., Cournac, A., Liti, G., Fischer, G., Zimmer, C., and Koszul, R. (2014b). Filling annotation gaps in yeast genomes using genome-wide contact maps. Bioinformatics 30, 2105–2113.

Meyer, F., Paarmann, D., D’ Souza, M., Olson, R., Glass, E.M., Kubal, M., Paczian, T., Rodriguez, A., Stevens, R., Wilke, A., et al. (2008). The metagenomics RAST server – a public resource for the automatic phylogenetic and functional analysis of metagenomes. BMC Bioinformatics 9, 386.

Peng, Y., Leung, H.C.M., Yiu, S.M., and Chin, F.Y.L. (2012). IDBA-UD: a de novo assembler for single-cell and metagenomic sequencing data with highly uneven depth. Bioinformatics 28, 1420–1428.

Selvaraj, S., Dixon, J.R., Bansal, V., and Ren, B. (2013). Whole-genome haplotype reconstruction using proximity-ligation and shotgun sequencing. Nat. Biotechnol. 31, 1111–1118.

Wood, D.E., and Salzberg, S.L. (2014). Kraken: ultrafast metagenomic sequence classification using exact alignments. Genome Biol. 15, R46.

